# Complete genome assemblies for three variants of the *Wolbachia* endosymbiont of *Drosophila melanogaster*

**DOI:** 10.1101/729061

**Authors:** Preston J. Basting, Casey M. Bergman

## Abstract

Here we report genome assemblies for three strains of *Wolbachia pipientis* assembled from unenriched, unfiltered long-read shotgun sequencing data of geographically distinct strains of *Drosophila melanogaster*. Our simple methodology can be applied to long-read datasets of other *Wolbachia*-infected species to produce complete assemblies for this important model symbiont.

---

W*olbachia pipientis* is a widespread bacterial endosymbiont that infects 40% of arthropod species (1) and induces a wide range of effects including cytoplasmic incompatibility, feminization, male-killing, and parthenogenesis (2). Currently, our understanding of the impact of *Wolbachia* on its hosts is limited by the lack of complete reference genomes for different *Wolbachia* strains, with only 18 of 84 *Wolbachia* assemblies in the NCBI assembly database as of Aug 2019 defined as complete.

Recently, Faddeeva-Vakhrusheva *et al.* (3) showed that a complete assembly of *Wolbachia* could be generated as a by-product of assembling the genome of a *Wolbachia*-infected arthropod species using PacBio long-read sequences. Based on this observation, we attempted to generate complete *Wolbachia* assemblies using long-read shotgun sequencing data for three geographically distinct *Drosophila melanogaster* lines (I23, N25, and ZH26) (4) that were previously identified as being infected by variants of the *Wolbachia* strain *w*Mel (5). To do this, all reads from PacBio whole genome shotgun sequences of each strain respectively were assembled using CANU v1.8 (6). Assemblies for each strain generated only one contig matching the *w*Mel reference genome (NC_002978.6) by BLASTN v2.9.0 (7), which in each case corresponded to the entire *Wolbachia* genome. The *Wolbachia* contig from each strain was circularized to match the *w*Mel reference genome using minimus2 from AMOS v3.1.0 (8), then polished using Arrow (SMRTlink v6.0.0.47841, Pacific Biosciences) and Pilon v1.23 (9) using Illumina reads from (5).

After polishing, we identified 54, 13, and 18 SNP/indel variants for the I23, N25, and ZH26 *Wolbachia* strains, respectively, relative to the the *w*Mel reference genome. The higher similarity of N25 and ZH26 *Wolbachia* strains and increased divergence of the I23 *Wolbachia* strain relative to the the *w*Mel reference genome is consistent with previous work showing that *Wolbachia* from N25 and ZH26 are both in clade III of the *w*Mel phylogeny (which also contains the *w*Mel reference genome), while *Wolbachia* from I23 is in clade I of the *w*Mel phylogeny (which is more divergent from the *w*Mel reference genome) (5, 10).

Our work extends that of Faddeeva-Vakhrusheva *et al.* (3) by showing that high-quality, complete genome assemblies of *Wolbachia* strains can be generated without experimental enrichment of symbiont DNA (e.g. (11, 12)). Successful *de novo* assembly of complete *Wolbachia* genomes directly from unenriched long-read sequences also demonstrates that it is unnecessary to computationally filter symbiont reads from host reads based on similarity to *Wolbachia* reference genomes prior to assembly (13, 14). As the cost of long-read sequencing decreases, we argue that direct sequencing and assembly of unenriched, unfiltered long-read datasets could be applied easily to other *Wolbachia*-infected arthropod and nematode species to expand the number of complete *Wolbachia* reference genomes.

**TABLE 1.** Accessions and statistics for assemblies produced and raw sequencing data used in this study.

| Assembly | Assembly Accession | Genome Size (bp) | GC (%) | PacBio read accession | # PacBio reads | Average PacBio read length (bp) | Illumina read accession | # Illumina reads | Avg Illumina read length (bp) |
| --- | --- | --- | --- | --- | --- | --- | --- | --- | --- |
| I23 | CP042444 | 1269137 | 35.24 | SRR7060336 | 31867 | 4140 | SRR1663552 | 547585 | 99 |
| N25 | CP042446 | 1267781 | 35.23 | SRR7060337 | 47126 | 4096 | SRR1663577 | 358462 | 98 |
| ZH26 | CP042445 | 1267436 | 35.23 | SRR7060339 | 47960 | 4407 | SRR1663599 | 364241 | 98 |

## Data availability

The assemblies produced in this study were deposited at NCBI under accession PRJNA557362. PacBio data used to generate these assemblies were published in (4) and are available under SRA accession SRP142531. Illumina data used to polish the assemblies were published in (5) and are available under SRA accession SRP050151.

## ACKNOWLEDGMENTS

We thank the Georgia Advanced Computing Resource Center for computational resources and Joshua Udall (Iowa State University) for providing access to raw PacBio Sequel data used in this project. This work was supported by a University of Georgia Research Education Award Traineeship (PJB) and the University of Georgia Research Foundation (CMB).

